# Systematic evaluation of chromatin immunoprecipitation sequencing to study histone occupancy in dormancy transitions of grapevine buds

**DOI:** 10.1101/2022.03.14.484366

**Authors:** Dina Hermawaty, Jonathan Cahn, Tinashe G. Chabikwa, Ryan Lister, Michael J. Considine

## Abstract

The regulation of DNA accessibility by histone modification has emerged as a paradigm of developmental and environmental programming. Chromatin immunoprecipitation followed by sequencing (ChIP-seq) is a versatile tool widely used to investigate *in vivo* protein-DNA interaction. The technique has been successfully demonstrated in several plant species and tissues; however, it has remained challenging in woody tissues. Here we developed a ChIP method specifically for mature dormant grapevine buds (*Vitis vinifera* cv. Cabernet Sauvignon). Each step of the protocol was systematically optimised, including crosslinking, chromatin extraction, sonication, and antibody validation. Analysis of histone H3-enriched DNA was performed to evaluate the success of the protocol and identify occupancy of histone H3 along grapevine bud chromatin. To our best knowledge, this is the first ChIP experiment protocol optimised for grapevine bud system.

## 2. Introduction

Chromatin immunoprecipitation (ChIP) enables the study of DNA-protein interactions and has become a method of choice for studying trans-regulation of gene expression, as well as post-translation histone modification. The technique was developed following a report demonstrated reversible crosslinking of nucleosome-DNA by formaldehyde (Jackson, 1978; Klockenbusch et al., 2012). In combination with several DNA assay techniques, such as southern blotting (Solomon et al., 1988 and Orlando et al., 1997), polymerase chain reaction (Hecht et al., 1996), microarray (Iyer et al., 2001), and sequencing (Johnson et al., 2007), the DNA sequence associated with the protein of interest may be identified. Forty years after its development, ChIP has been extensively used to study epigenetic regulation in animal and yeast cells, but only recently applied in plants (Johnson et al., 2001 and Wang et al., 2002). The delay in uptake of ChIP in plant science was due to several impediments, particularly: (1) a large amount of tissue is typically needed, (2) the presence of cell walls required vigorous physical disruption therefore sample loss during the process is unavoidable and resulted in low DNA yield, (3) co-extraction and precipitation of interfering compounds often problematic for downstream analysis such as PCR/ qPCR and library preparation, (4) limited availability of ChIP-grade antibodies specific for plant cells often leading to a false-negative signal, and (5) the comprehensive ENCODE guidelines for model biological system is not always applicable for plant biology research.

The intriguing and complex regulation of plant developmental processes, as a response to environmental stimuli, has driven many studies on gene expression regulation in an epigenetic context. The vernalisation requirement for flowering of Arabidopsis is established by the flowering repressor FLOWERING LOCUS C (FLC), whereby chilling-dependent histone modification of the FLC locus represses transcription and hence enables flowering (Michael and Amasino, 1999; Halliwell et al., 2006). As histones are widely conserved and several commercial antibodies available, ChIP has been successfully applied to non-model plant studies also, including dormancy in perennial buds (Leida et al., 2012; Saito et al., 2015; and de la Fuente et al., 2015). To date, protocols guiding ChIP experiments in plant systems, such as Arabidopsis (Saleh et al., 2008), tomato (Ricardi et al., 2010), maize (Haring et al., 2007) followed by DNA microarray hybridization (Reimer and Turck, 2010) or sequencing (Kaufmann et al., 2010) have been published. However, the variables amongst these studies illustrate the need to tailor conditions to each experiment, and in particular each tissue type (Park, 2009; Landt et al., 2012). As such, protocols established for soft tissues such as leaves (Saleh et al., 2008) or seedlings (Ricardi et al., 2010) are likely to be ineffective for seed (Haque et al., 2018) or wood forming tissues (Li et al., 2014a). Further, metastudies have shown that even commercially available ChIP-grade antibodies may fail control tests for specificity (Egelhofer et al., 2011). In some cases, batch information of these validation steps is available either on the ENCODE Project website (Davis et al., 2017) or subsites (Egelhofer et al., 2011) or via the manufacturer. Alternatively, the antibody/s must be validated before commencing ChIP experiment (Landt et al., 2012). Procedures and criteria for antibody validation have been well-outlined by members of the ENCODE Project, however these were specifically developed for animal tissues, and hence neglect for example the additional constraints of working with plant cell walls and particularly lignified tissues.

The ChIP workflow is summarised in **Figure 1**. In brief, the interaction of protein and DNA (collectively known as chromatin) is crosslinked in vivo by incubation of tissue in formaldehyde solution. The crosslinked chromatin is then fragmented by sonication which breaks the chromatin into short fragments that are suitable for the subsequent processes. The protein-DNA complex is co-precipitated using antibody allowing selective precipitation of DNA that interacts with protein of interest. The precipitated DNA is released from the protein by reverse crosslinking and subsequently assayed to identify the sequence. Each step in the ChIP procedure is prone to high variability; for example, sonication must be titrated to ensure the optimal size of chromatin while preventing damage. Similarly, for crosslinking, insufficient crosslinking could cause poor preservation of chromatin and its associated protein and significantly reduce the yield of DNA at the end of the immunoprecipitation process (Orlando, 2000). Alternatively, excessive crosslinking can make the chromatin brittle and prevent efficient reversibility of the crosslinking at subsequent steps. Therefore, optimisation needs to be systematic in order that the method is robust and reproducible, yielding maximum enriched-DNA (**Figure 1**, arrow).

**Figure 1.**
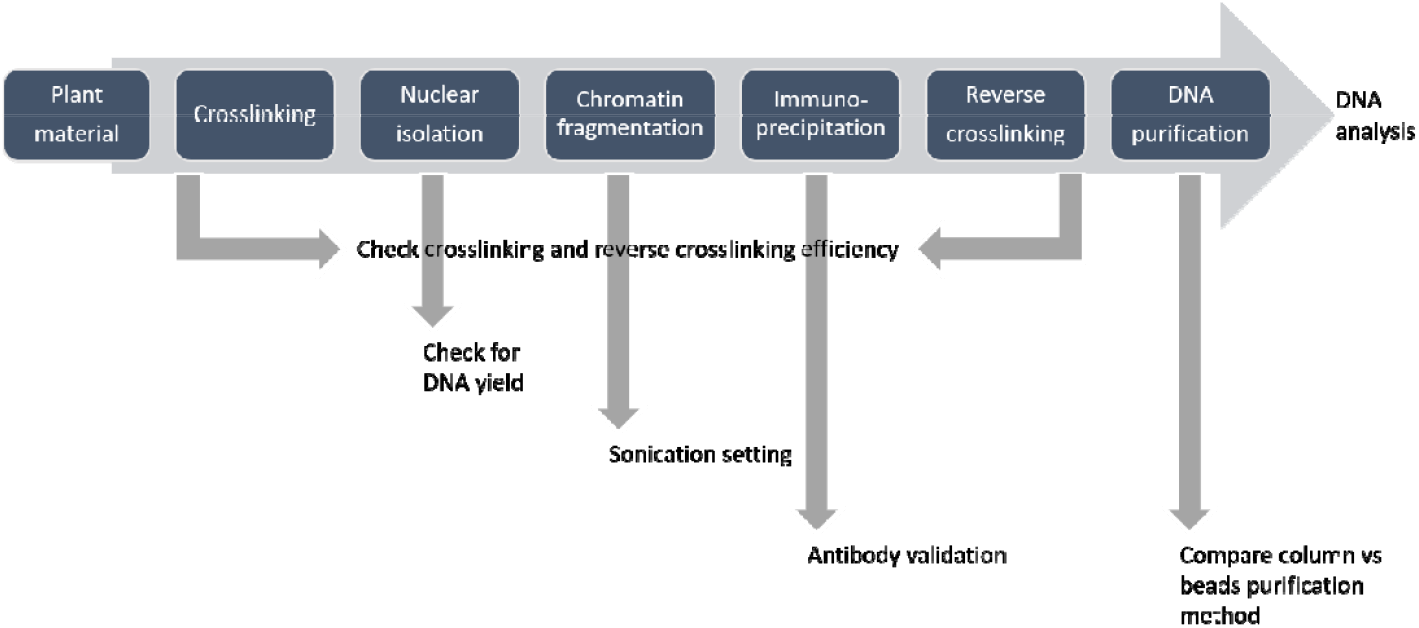
Chromatin immunoprecipitation workflow with checkpoints indicated by the grey arrow.

The ChIP protocol we describe is a modified procedure from a protocol optimised for wood-forming xylem tissue developed by Li et al. (2014) which provides a guide to cope with the difficulties of working with woody tissue. Systematic optimisation was performed according to ENCODE guidelines for ChIP experiment (Landt et al., 2012.) and other recommendations from previously published ChIP protocols with plant tissue (Ricardi et al., 2010; Haring et al., 2007; Song et al., 2016). Chromatin immunoprecipitation was performed using a ChIP kit manufactured by Abcam to eliminate washing steps after immunoprecipitation which often contribute to loss of enriched-DNA. Finally, we performed, DNA sequencing and identified gene that was occupied by histone H3 protein.

## 3. Materials and Equipment

### Plant material and treatment

The mature dormant buds of *Vitis vinifera* (L.) cv. Cabernet Sauvignon (N+2, Lavee and May, 1997) were collected from a vineyard in Margaret River, Australia (34 °S, 115 °E) at three time points; March, May, and August. Each cutting was consisting of 4 mature buds from node 4 to 7. The canes were immediately transported to the lab in damp newsprint in an insulated box and stored at 22 °C for up to 24 hours. Treatment with hydrogen cyanamide (H_2_CN_2_; Sigma-Aldrich #187364) was done by submerging the node into 1.25 % (w/v) [300 mM] H_2_CN_2_ for 30 seconds. Control buds were treated in the same manner with water (W). The explants were then stored in the dark for 24 hours at room temperature before being crosslinked.

### Reagents

- Sucrose (Chem-Supply, Australia, cat. no. SA030-500G)
- UltraPure 1 M Tris-Cl pH8 (Invitrogen, Australia, cat. no. 15568-025)
- 0.5 M EDTA pH 8 (Invitrogen, cat. no. AM9260G)
- Paraformaldehyde (Sigma-Aldrich, Australia, cat. no. P6148-500G)
- Glycine (Chem-Supply, cat. no. GA007-500G)
- β-mercaptoethanol (Sigma-Aldrich, cat. no. 63689-100ML-F)
- Polyvinylpyrrolidone (Sigma-Aldrich, cat. no. PVP40-100G)
- Triton X-100 (Sigma-Aldrich, cat. no. T9284-100ML)
- NaCl (Sigma-Aldrich, cat. no. S7653-1KG)
- Sodium dodecyl sulfate (SDS, Merck, Australia, cat. no. 8.17034.1000)
- Hydrochloric acid (Sigma-Aldrich, cat. no. 320331-2.5L)
- Miracloth (Merck-Millipore, Australia, cat. no. 475855)
- UltraPure phenol:chloroform:isoamyl alcohol 25:24:1 (v/v) (Invitrogen, cat. no. 15593031)
- Ethidium bromide (Sigma-Aldrich, cat. no. E1510)
- Absolute ethanol (Merck, cat. no. 1.00983.2511)
- Agarose (Thermo Scientific, Australia, cat. no. 16500100)
- 1 kb DNA ladder (Promega, USA, cat. no. G5711)
- Mini-PROTEAN TGX (Tris-Glycine eXtended), 4-15% precast gradient polyacrylamide gel (Biorad, Australia, cat. no. 161-1104EDU, 10-well, 30 µl, 8 × 10 cm (W x L))
- 10x Tris/Glycine/SDS Buffer (Biorad, cat. no. 161-0732)
- 4X Laemmli buffer (Biorad, cat. no. 1610747)
- Protein marker (Blue Star Pre-stained Protein Marker, Nippon Genetics, Japan, cat. no. MWP03)
- Immun-Blot^®^ PVDF membrane, precut, 7 × 8.4 cm (Biorad, cat. no. 1620174)
- Extra thick blot filter paper, precut, 8 × 13.5 cm (Biorad, cat. no. 1703966)
- Trans-Blot^®^ SD Semi-Dry Electrophoretic Transfer Cell (Biorad, cat. no. 1703940)
- Primary antibodies: Histone H3 – nuclear loading control rabbit pAb (Abcam, Australia, cat. no. ab1791), Histone H3K4me3 antibody rabbit pAb (Active Motif cat. no. 39915), Histone H3K27me3 antibody rabbit pAb (Active Motif cat. no. 39155)
- Goat anti-rabbit IgG HRP conjugated secondary antibody (Santa Cruz Biotechnology, cat. no. SCZSC-2030)
- Pierce™ ECL Western Blotting Substrate (Thermo Scientific, Australia, cat. no. 32109)
- ChIP kit plant (Abcam, cat. no. ab117137)
- NEBNext^®^ Ultra™ II DNA Library Prep Kit for Illumina^®^ (New England Biolabs, cat. no. NEB.E7645G).
- NEBNext^®^ Multiplex Oligos for Illumina^®^ (New England Biolabs, cat. no. NEB.E7335G).
- Agentcourt AMPure XP beads (Beckman Coulter Life Science, USA, cat. no. A63881)
- 4’,6-Diamidino-2-Phenylindole dihydrochloride (DAPI, Sigma, cat. no. 102M4012V)

### Equipment

- Vacuum chamber
- Vacuum pump
- Aluminium foil
- Conical tubes (50 mL and 15 mL)
- Mortar and pestle
- Rotator
- Vortex (Velp Scientifica, Italy)
- ULTRA-TURRAX homogeniser (model T25 basic, IKA, Germany)
- Refrigerated centrifuge (model 5810R, Eppendorf)
- Fix-angle rotor (model F45-30-11 and F34-6-38, Eppendorf)
- Microcentrifuge tube (1.5 and 2 mL)
- Focus-ultrasonicator (model S220, Covaris, USA)
- miliTUBE 1 mL AFA fibre (Covaris, cat. no. 520130)
- Hot water bath (model B-491, Buchi, Switzerland)
- NanoDrop (model ND-1000, Thermo Fischer Scientific, Australia)
- Qubit fluorometer (model Qubit 3.0, Thermo Fischer Scientific, Australia)
- Bioanalyzer (Agilent 2100 bioanalyzer, Agilent, Australia)
- Electrophoresis system (Mini Gel II, Select BioProduct, USA)
- Mini-PROTEAN Electrophoresis system (Biorad)
- ChemiDoc MP system (Biorad)
- DynaMag™-2 Magnet (Thermofischer scientific, cat. no. 12321D)
- Axioscope optical microscope (Zeiss, Oberkochen, Germany) equipped with plan-neofluar objectives, UV or blue epi-illumination and differential interference contrast filters.
- Axiocam digital camera (Zeiss Oberkochen, Germany)

### Reagent setup

- **Formaldehyde (16%)** Dissolve 4 grams of paraformaldehyde in 21 mL of water and add 1µL NaOH (10 M). Stir and heat (no more than 68 °C) until in solution. Let cool to room temperature and bring the solution to a final volume of 25 mL.
- **Sucrose, 2M** Dissolve 68.46 grams of sucrose in 56 mL water. Stir and heat until in solution and bring to a final volume of 100 mL. Freshly prepare the solution prior to experiment.
- **Glycine, 2M** Dissolve 15 grams of glycine in 80 mL of water. Stir until in solution and bring to a final volume of 100 mL. Store solution at 4 °C and allow solution to reach room temperature (RT) before use.
- **10X Protease Inhibitor** Dissolve cOmplete protease inhibitor, EDTA-free in 5 mL water or dissolve cOmplete protease inhibitor, mini-tablet, EDTA-free in 1 mL water. Vortex until in suspension. Freshly prepare the suspension prior to experiment. Keep at 4 °C.
- **Triton X-100, 10% (v/v)** Dissolve 5 mL of Triton X-100 in 40 mL water. Stir slowly until in solution and bring to a final volume of 50 mL. Store at Store solution at 4 °C.
- **NaCl, 5 M** Dissolve 29.22 grams of NaCl in 80 mL water. Stir until in solution and bring to a final volume of 100 mL. Autoclave and store solution at RT.
- **SDS, 10% (w/v)** Dissolve 10 grams of SDS in 80 mL water. Stir slowly and heat until in solution. Bring the solution to a final volume of 100 mL. Autoclave and store solution at RT.
- **Buffer 1** contains 0.4 M sucrose, 10 mM Tris-Cl, 2.5% (w/v) PVP-40, 5 mM β-mercaptoethanol, 1× Roche cOmplete protease inhibitor, EDTA-free. Freshly prepare the buffer prior to experiment. Pre-chilled before use. Add β-mercaptoethanol and protease inhibitor to the buffer before use.
- **Buffer 2** contains 0.25 M sucrose, 10 mM Tris-Cl, 10 mM MgCl2, 1% (v/v) Triton X-100, 5 mM β-mercaptoethanol, 1× Roche cOmplete protease inhibitor, EDTA-free. Freshly prepare the buffer prior to experiment. Pre-chilled before use. Add β-mercaptoethanol and protease inhibitor to the buffer before use.
- **Buffer 3** contains 1.7 M sucrose, 10 mM Tris-Cl, 0.15% (v/v) Triton X-100, 5 mM β-mercaptoethanol, 1× Roche cOmplete protease inhibitor, EDTA-free. Freshly prepare the buffer prior to experiment. Pre-chilled before use. Add β-mercaptoethanol and protease inhibitor to the buffer before use.
- **Lysis buffer** contains 50 mM Tris-Cl, 10 mM EDTA, 0.1% (v/v) SDS, 1× Roche cOmplete protease inhibitor, EDTA-free. Freshly prepare the buffer prior to experiment. Pre-chilled before use. Add protease inhibitor to the buffer before use.
- **Ethanol, 70% (v/v)** add 30 mL of water into 70 mL of absolute ethanol. Prepare solution prior to experiment.
- **Tris-EDTA buffer with low EDTA (TE-lowE)** TE-lowE contains 10 mM of Tris-Cl and 0.1 mM EDTA pH.8. Store solution at 4 °C.
- **Transfer buffer** Transfer buffer contains 39 mM glycine, 48 mM tris base, 0.05%(v/v) SDS, 20% (v/v) methanol. Adjust pH to 8.3 and store at 4 °C.
- **Tris-buffered saline (TBS) 10X** Dissolve 24.23 grams of Tris base and 80.06 grams of NaCl in 800 mL water. Stir until in solution and adjust pH to 7.6 with HCl. Bring the solution to a final concentration of 1 L.
- **Tris-buffered saline with tween (TBST)** TBST contains 1X TBS, 0.5% (v/v) Tween-20. Stir slowly. Store buffer at 4 °C.
- **Blocking buffer** Dissolve 5% (w/v) non-fat milk in TBST. Stir until in suspension and keep at RT. Prepare buffer prior to experiment.
- **DAPI, 1 mg/mL** Dissolve 1 mg of DAPI dye in 1 mL water. Vortex until in solution. Keep in dark at 4 °C

### Procedure

#### Tissue collection

1. The buds from node 4 to 7 were excised from the cane and dissected into half, longitudinally, to increase surface exposed to crosslinking buffer then immediately immersed in a fixative solution. We used whole buds in this experiment for the convenience of bud harvesting.

#### Crosslinking

2. Immediately put the bud into conical tube contains 25 mL **CROSSLINKING BUFFER**, repeat this until 100 buds are obtained (ca. 2.5 grams). Crosslink the buds for a total of 15 minutes under cycled vacuum infiltration (5 min/ release/ mix, repeat three times) at room temperature. **NOTE:** Excessive exposure to crosslinking agents may result in inefficient DNA fragmentation and protein denaturation. Since buds need to be excised from the canes for this experiment, it took some time to harvest 5-10 grams of buds. We suggest cutting as many buds as possible in 30 minutes then immediately proceed with vacuum infiltration. In our case, we handled 100 buds at a time.
3. Quench the crosslinking reaction by addition of 2 M glycine to a final concentration of 200 mM, followed by 5 minutes cycled vacuum.
4. Rinse crosslinked tissue with water twice. Dry tissue between absorbent paper then put them on the foil.
5. Snap freeze tissue in liquid nitrogen and store at −80 °C until required.

#### Nuclear isolation

6. Unless otherwise indicated, all step must be performed at 4 °C, and the sample must be kept on ice all the time.
7. Grind crosslinked buds to a fine powder in liquid nitrogen using mortar and pestle. Always grind a small amount of tissue at a time, then collect powder into a new 50 mL conical tube. Repeat grinding until all 10 grams of crosslinked buds are ground. The conical tube must be kept on dry-ice all the time. **NOTE**: one 50 mL canonical tube is suitable for 5 grams of tissue powder. When working with 10 grams tissue, split the ground powder into two new tubes.
8. Mix the powder with seven volumes of **BUFFER 1** in 50 mL conical tube (e.g. 35 mL for every 5 grams tissue). Homogenize using a vortex and an ULTRA-TURRAX homogenizer at 9000 rpm for 15 seconds. Further mix suspension in rotating wheel for 20 minutes at 4 °C. **NOTE:** Complete homogenisation is important to get a maximum DNA yield. **CHECKPOINT:** Comparing DNA yield obtained from vortex homogenization vs ULTRA-TURRAX may be needed to optimise the homogenisation method.
9. Pass the mixture through three layers of Miracloth saturated with Buffer 1 into new 50 mL conical tube. Squeeze the Miracloth to collect all the liquid.
10. Centrifuge suspension at 2,880 *g* for 10 minutes at 4 °C. Discard supernatant.
11. Gently resuspend pellet in 2 mL of **BUFFER 2** and transfer suspension into a new 2 mL microcentrifuge tube.
12. Centrifuge suspension at 12,000 *g* for 10 minutes at 4 °C. Discard supernatant.
13. Repeat step 10 to 12 once.
14. Gently resuspend pellet in 500 µL of **BUFFER 3**. Carefully layer the suspension on top of 1.5 mL cushion of **BUFFER 3** in a new 2 mL microcentrifuge tube. **NOTE**: Pellet may be difficult to resuspend. A disposable tissue grinder pestle can be used to carefully loosen the pellet followed by pipetting up and down.
15. Centrifuge sample at 16,000 *g* for 60 minutes at 4 °C. Discard supernatant.
16. Gently resuspend pellet in 700 µL of **LYSIS BUFFER**. Take 50 µL for the no-sonication control and keep the resuspended pellet on ice. **CHECKPOINT:** check yield of DNA and validate antibody (**Supplementary Information S1** and **S2**). **CHECKPOINT:** Nuclei integrity can be checked by adding DAPI dye to a final concentration of 10 mg/mL and examine nuclei using an epiluminescence microscope (**Figure 6**).

#### DNA fragmentation

17. Transfer nuclei suspension into miliTUBE being sure to fill the tubes with lysis buffer (a little more than 1 mL per tube).
18. Sonicate the DNA in Covaris S220 focus-ultrasonicator for 12 minutes following manufacture’s setting for high cell chromatin shearing, i.e. 5 % Duty Cycle, 4 intensity, 140 W peak incident power, 200 cycles per burst, 6 °C bath temperature, frequency sweeping power mode, continuous degassing mode, and level 8 water. Transfer sonicated DNA into a new 1.5 mL. **CHECKPOINT**: Take 50 µL aliquots after 6, 8, and 10 minutes to compare DNA fragmentation and each time replace with the same amount of lysis buffer. Keep sample on ice.
19. Centrifuge sonicated and non-sonicated DNA at 16,000 *g* for 10 minutes at 4 °C. Transfer clean supernatant into a new 1.5 mL microcentrifuge tube.
20. Proceed immediately to step 21 for chromatin immunoprecipitation. DNA can be stored at – 20 °C and proceed to **Supplementary Information S1** for DNA fragmentation efficiency examination.

#### Chromatin immunoprecipitation and reverse crosslinking

The following chromatin immunoprecipitation and reverse crosslinking procedure are adapted from ChIP kit plant from Abcam with some modification.

21. Determine the number of strip wells required. Leave these strips in the plate frame (remaining unused strips can be placed back in the bag. Seal the bag tightly and store at 4 °C).
22. Wash strip wells once with **150 μL** of **WASH BUFFER**.
23. Add 100 μL of the **ANTIBODY BUFFER** to each well and then add the antibodies:
  - **3 μg** of an antibody of interest (H3K27me3 and H3K4me3).
  - **2 μg** of anti-histone H3 as a positive control.
  **NOTE:** ChIP typically requires 1-10 µg per ChIP reaction. Optimising the amount used per reaction is a further variable to consider, however here the amount chosen followed manufacturer recommendations. In our experiment with grapevine buds, three reactions (wells) were prepared for each histone H3 modified antibody and two reactions for histone H3 antibody (**Figure 2)**.

**Figure 2.**
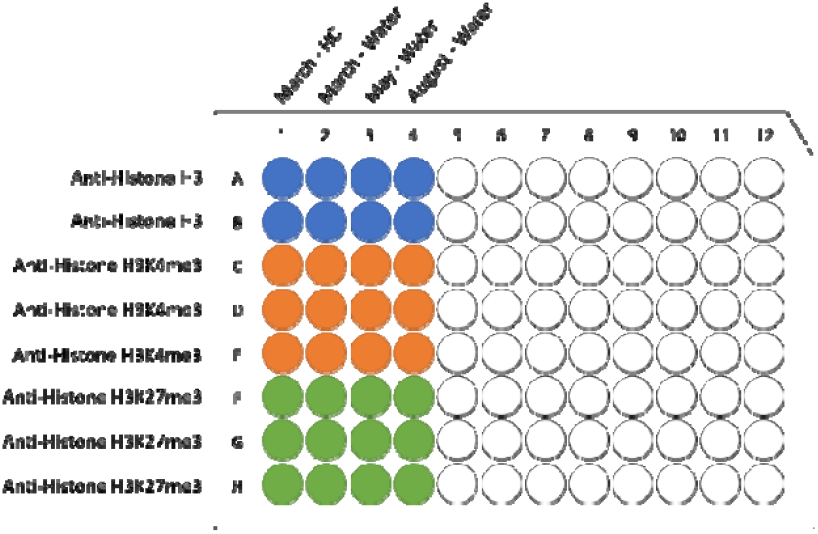
ChIP assay plate map. Incubation of chromatin and antibodies is done in an assay-well provide in Abcam’s ChIP kit plant. Each well is designed for one ChIP reaction using 100 µL fragmented DNA. In our experiment, multiple wells were used per antibodies, i.e. 2-well for anti-histone H3 (blue) and 3-well each for anti-Histone H3K4me3 (orange) and anti-Histon H3K27me3 (green), with each column represent different sample.
24. Cover the strip wells with **Parafilm M** and incubate at room temperature for 90 minutes.
25. After incubation, remove the incubated antibody solution and wash the strip wells **three times** with **150 μL** of the **ANTIBODY BUFFER** by pipetting in and out.
26. Remove **15 μL** of chromatin aside to a 0.5 mL vial. Label the vial as **“input DNA”** and then place on ice. **NOTE:** the amount of input DNA is 5 % from the total volume of chromatin used per histone H3 modifies antibodies, i.e. 5 % from 300 µL.
27. Transfer **100 μL** of **chromatin from step 19** to each antibody-bound strip well. Two and three reactions (wells) are used for Histone H3 and Histone H3-modified immunoprecipitation. **NOTE:** Concentration of SDS in LYSIS BUFFER (step 15) is 0.1 %; therefore, no sample dilution needed.
28. Cover the strip wells with **Parafilm M** and incubate at 4 °C for overnight on an orbital shaker (50-100 rpm).
29. Remove supernatant. Wash the wells six times with **150 μL** of the **WASH BUFFER**. Allow 2 minutes on a rocking platform (100 rpm) for each wash.
30. Wash the wells once (for 2 minutes) with **150 μL of 1X TElowE BUFFER**.
31. Add **40 μL of the DNA Release mix**, containing 1 µL Proteinase K (10 mg/mL) and 40 µL DNA RELEASE BUFFER, to the samples (including the “input DNA” vial).
32. Cover the sample wells with strip caps and incubate at 65 °C in a water bath for 15 minutes. Following incubation at 65 °C do a quick spin to collect all suspension at the bottom of the plate.
33. Add **40 μL of the REVERSE BUFFER** to the samples and to a vial labelled as “input DNA”; mix and re-cover the wells with strip caps and incubate at 65 °C in a water bath for 90 minutes. Quick spin plate at RT.
34. Combine solution from the same histone antibody (2 wells for Histone H3 and 3 wells for Histone H3 modified).

#### DNA purification with AMPure Beads

35. Add 1.8X volume of AMPure XP beads to IP enriched and input DNA. **NOTE**: This step will bind DNA fragments size from 100 bp and larger.
36. Mix reagent and sample thoroughly by **pipette mixing ten times**.
37. Let mixed samples incubate for 15 minutes at room temperature for maximum recovery. **NOTE**: pipette mixing is preferable to vortexing as it tends to be more reproducible. The colour of the mixture should appear homogenous after mixing.
38. Place on a magnetic rack for 5 minutes (wait for solution to clear before proceeding to the next step).
39. With tube still in the magnetic rack, aspirate the clear solution from tube and discard.
40. Keep the sample in magnetic rack and add 1 mL of freshly prepared 70 % ethanol.
41. Incubate for 30 seconds at room temperature. Aspirate out the ethanol and discard.
42. Repeat ethanol wash one more time.
43. Illumina recommended at least 15 minutes drying time but longer drying time may be required. **NOTE:** ensure all traces of ethanol are removed but avoid over-drying the beads, which will significantly decrease elution efficiency (beads will appear cracked if over dried).
44. Remove the tube from the magnetic rack, add 10 μL TElowE and pipet up and down several times until pellet beads are completely resuspended. **NOTE:** Standard TE **must not be used** at this step.
45. Incubate at room temperature for 2 minutes. Place in the magnetic rack for 5 minutes.
46. Transfer 9 μL of the supernatant to a 0.2 mL PCR tube.
47. Repeat step 44-46 once. DNA is now ready for use or store at – 20 °C.

### Sequencing

The library was constructed using NEBNext^®^ Ultra™ II DNA Library Prep Kit for Illumina^®^ following manufacturer’s low-input ChIP-seq protocol (**Supplementary Information S3**). The library for input and histone H3-enriched DNA each from March (water- and H_2_CN_2_-treated), May and August samples were sequenced at Genewiz Genomics Centre (Suzhou, China) as pair-end (PE), 150 bp for an average of 40 million of reads per sample. Raw reads were trimmed for quality and adaptors using Trimmomatic v0.39 (Bolger et al., 2014). Post-trimming read quality was assessed using FastQC and results were aggregated using MultiQC (Ewels et al., 2016). The remaining reads were mapped to the 12X V1 *Vitis vinifera* PN40024 reference genome (Jaillon et al., 2007) using the Burrows-Wheeler Aligner (BWA) (Li et al., 2009). Peak calling was conducted using MACS2 software version 2.1.0 (https://github.com/taoliu/MACS) with cut off q-value < 0.05. The annotatePeaks.pl algorithm of the HOMER (Hypergeometric Optimization of Motif EnRichment) suite of tools (Heinz et al., 2010) was used to annotate the peaks. DeepTools (Ramírez et al., 2016) was used to process the mapped reads data for creating normalized coverage files in standard bedGraph and bigWig file formats for visualisation and comparison between different files. Functional category enrichment was performed for genes that were enriched by histone H3 using topGO package following a grapevine-specific functional classification of 12X V1 predicted transcript (Grimplet et al., 2012) with modification according to the GO database (Ashburner et al., 2000). A Fisher’s exact test (*P* < 0.05) was carried out in topGO to compare each study list with the list of total non-redundant transcript housed in grapevine 12X V1 gene predictions (Grimplet et al., 2012). The gene ontology GO terms were further simplified using REVIGO allowing similarity of 0.5 (Supek et al., 2011).

## Results

### Crosslinking by vacuum infiltration

Infiltration with 15 minutes cycled vacuum (5 min vacuum/release/mix × 3) and without vacuum was compared to determine a suitable infiltration method for grapevine buds. Complete infiltration was indicated by the movement of buds to the bottom of the tube as the buds’ density become higher after infiltration of crosslinking buffer (**Figure 3**).

**Figure 3.**
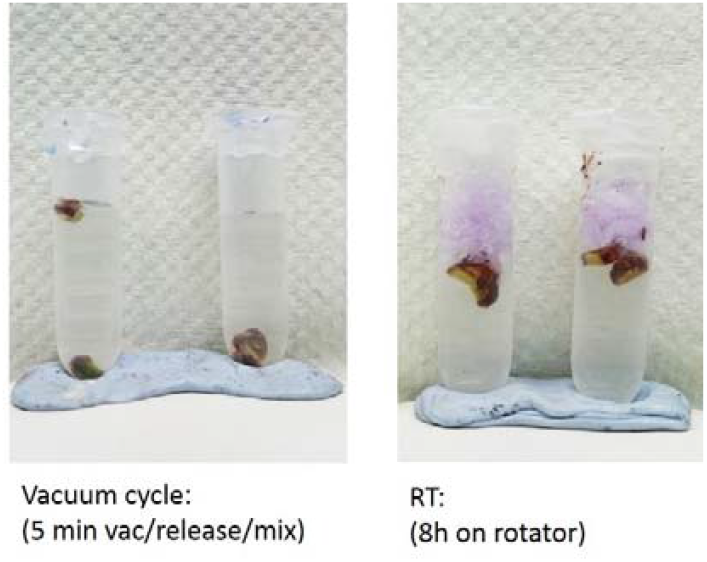
Fixative infiltration optimisation. Buds were cut into half before immersed into the fixative solution. **Left**: cycled vacuum was applied by performing three cycles of 5 minutes vacuum, release, and mix at room temperature. **Right**: a stopper (light purple) was placed on top of the solution to keep the sample submerged in the fixative solution the tubes were left on a rotator for 8 hour at room temperature. An efficient penetration of the fixative was evident after vacuum indicates by increasing of the bud density which causes buds sunk into the bottom of the tube. Cycled vacuum method also allows short crosslinking process which is preferred for ChIP analysis.

The phenol:chloroform:isoamyl alcohol (PCI) solution separates nucleic acid and protein based on its solubility in the solvents, i.e. nuclei acid is soluble in aqueous phase while protein in organic phase. Excessive crosslinking or ineffective reverse crosslinking will retain interaction between DNA and protein and therefore reduce the amount of DNA in the aqueous phase because the protein-DNA complex will be soluble in the organic phase instead. Crosslinking efficiency of our protocol was then assessed by comparing amount of DNA in the aqueous phase from crosslinked and non-crosslinked bud, treated with or without reverse crosslinking. In non-crosslinked bud (**Figure 4, lane 1-3**), DNA was soluble in the aqueous phase with or without reverse crosslinking treatment. In contrast, when crosslinking was performed, DNA can only be recovered from the aqueous phase if reverse crosslinking procedure was conducted (**Figure 4, lane 6**). The overnight reverse crosslinking procedure can be done as an alternative to a shorter duration without affecting DNA recovery (**Figure 4, lane 7**). Absence of DNA at **lane 5** confirmed the successful crosslinking procedure which maintains the protein-DNA interaction, while presence of DNA at **lane 6-7** demonstrates efficiency of our crosslinking allowing release of DNA from protein.

**Figure 4.**
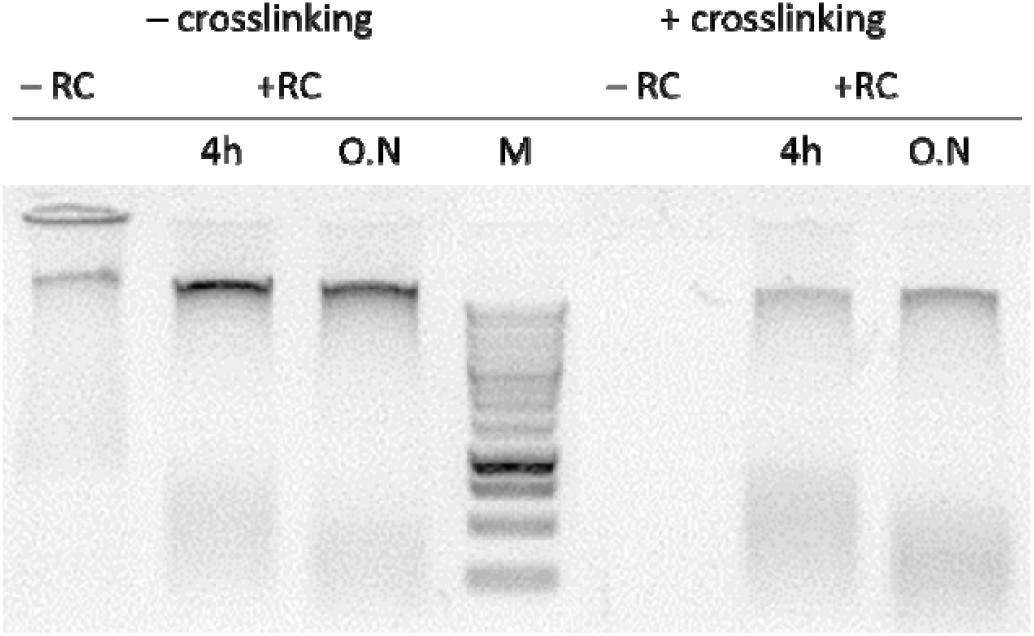
Crosslinking and reverse crosslinking efficiency. Nuclear extract was prepared from grapevine buds without **(–) crosslinking** and with **(+) crosslinking** treatment. Grapevine buds were crosslinked in crosslinking buffer containing 1% formaldehyde for 15 minutes (3×5 minutes vacuum cycles) at room temperature. The sample was **reverse crosslinking (+RC)** for 4 hour and over the night (O/N) or **not (–RC**). DNA was purified using phenol/chloroform extraction followed by ethanol precipitation. DNA recovery was compared between samples with and without crosslinking.

### Chromatin yield and nuclei integrity

Disruption of antigen-antibody interaction mainly avoided in most ChIP protocols by using 1 % SDS in lysis buffer and further dilute the chromatin suspension after DNA fragmentation to reduce the SDS concentration to 0.1%. We obtained the highest DNA yield using 1 % SDS (**Figure 5, lane 3-4**); however, a considerable increase of DNA yield was observed after application of 6 minutes of sonication in sample lysed using low detergent concentration (**Figure 5, lane 1-2 and 5-6**. An aliquot of six minutes sonicated nuclei suspension (**see procedure, step 16**) was stained with DAPI (1 µg/ mL) and subjected to microscopic observation to assess integrity of the nuclei. The micrograph showed a uniform, intact and well-separated nucleus (**Figure 6)**.

**Figure 5.**
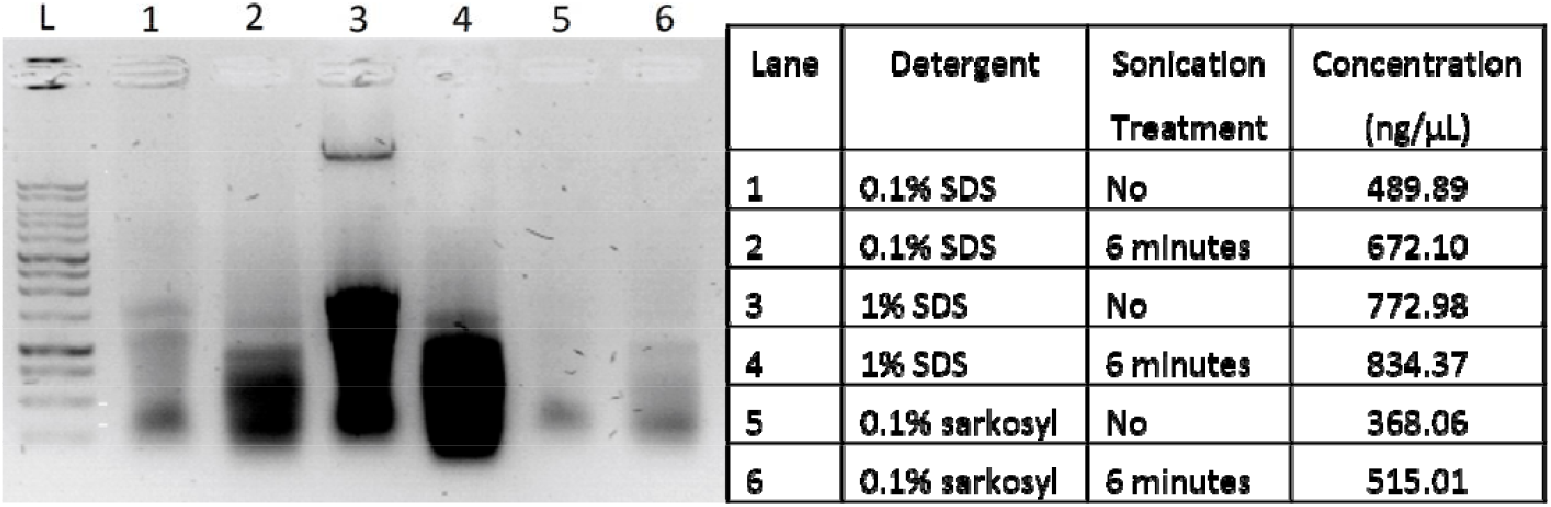
The combination of type and concentration of detergent in the lysis buffer and application of sonication resulted in a different yield of DNA. L: 1Kb DNA ladder (Promega #G5711) in 1% agarose gel, DNA quantification was performed using a NanoDrop 1000.

**Figure 6.**
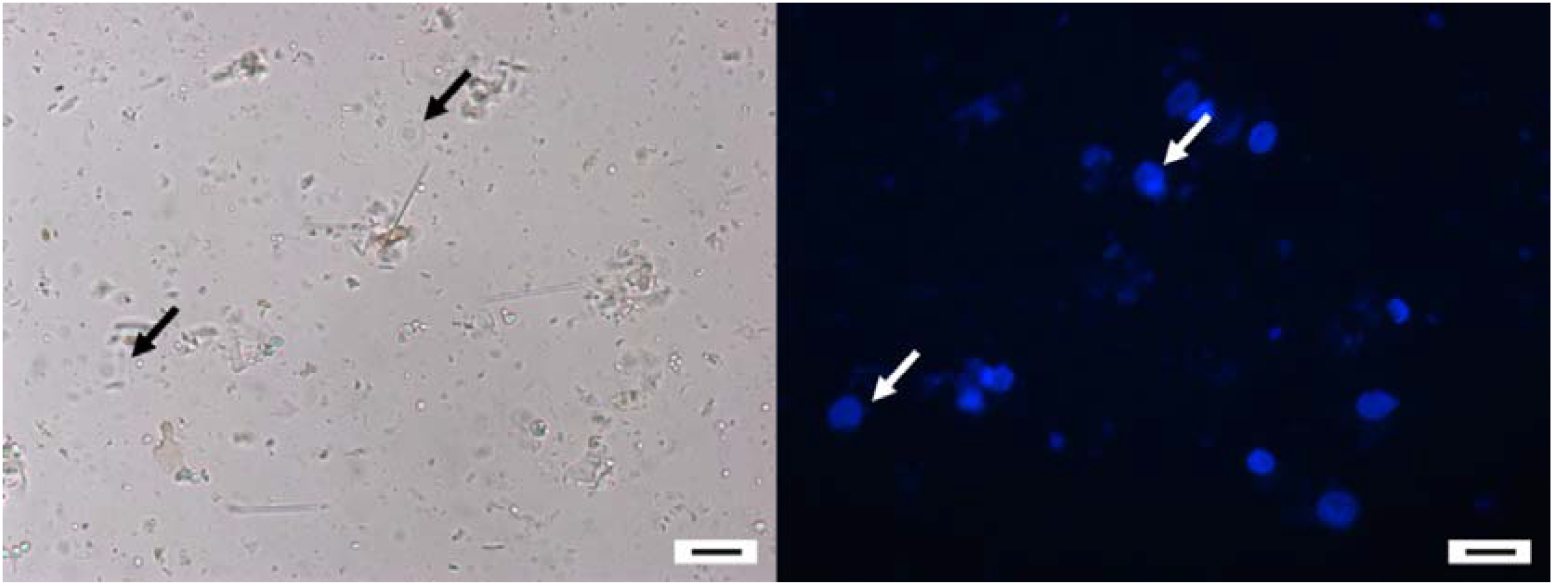
Nuclei integrity assessment by examination under a microscope. DAPI stain DNA specifically at the A-T rich region and will emit blue fluorescence light which can be observed using an epiluminescence microscope. The image was taken using DAPI filter (exciter filter BP 365/12, chromatic beam splitter FT 395, and barrier filter LP 397). Bar = 5 µm

### DNA fragmentation

A sonicator setting to produce an average of 300 bp fragment was used, following the default setting provided by Covaris S220 Focussed-ultrasonicator manufacture. In general, short DNA fragments were gradually accumulated as sonication duration increased (**Figure 7**). After 8 minutes of sonication, the average fragment size was not changed coincided with an increase of fragment within 200-400 bp range. Increasing the duration of sonication to 10 minutes, the accumulation of DNA fragments in the 200-400 bp range increased without causing further fragmentation of the short DNA.

**Figure 7.**
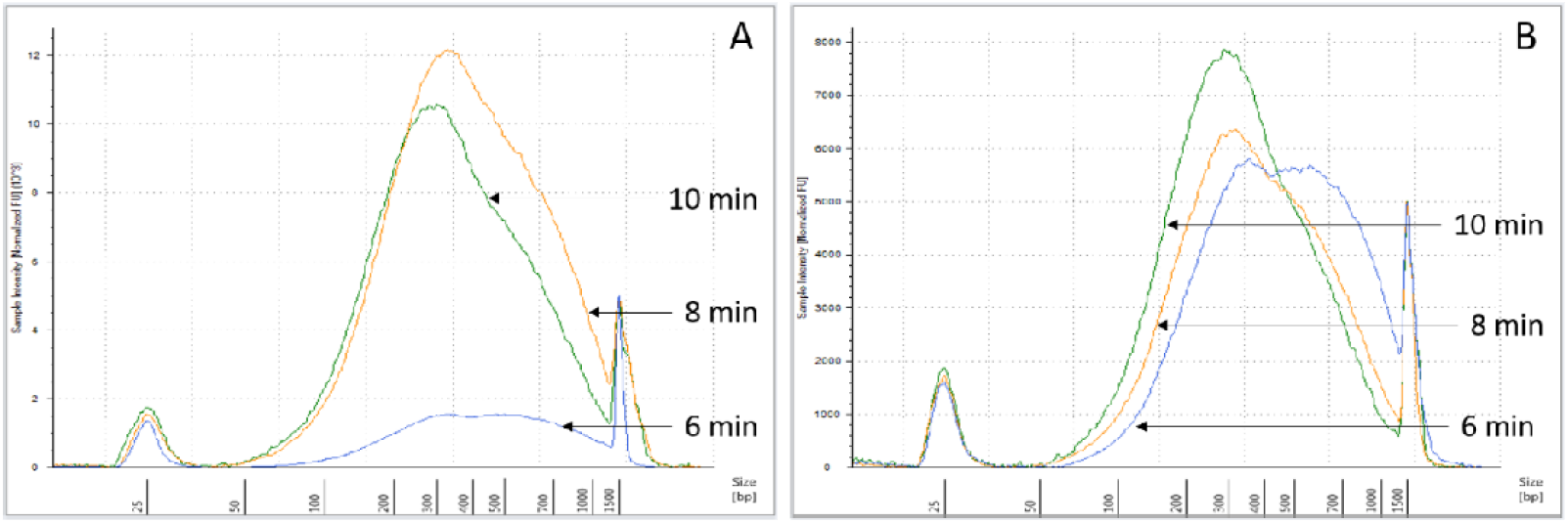
Optimisation of chromatin fragmentation. Chromatin fragmentation was optimised to obtain suitable DNA fragment size for ChIP-seq, i.e. 200-400 bp. Chromatin extracted using 0.1% (**A**) and 1% SDS (**B**) were sonicated for 6 (blue), 8 (yellow) and 10 (green) minutes. Distribution of DNA fragment size was analysed using Agilent Bioanalyzer. Accumulation of smaller DNA fragment was linear to sonication duration with suitable average fragment size was obtained after 8 minutes, and more accumulation of fragment size from 200-400 bp observed after 10 minutes sonication.

### Yield of immunoprecipitated-DNA

Three different methods to purify the immunoprecipitated-DNA were tested in which the lowest DNA recovery was produced by column purification method while the paramagnetic beads (AMPure XP) resulted the highest DNA yield (**Table 1**). Therefore, we substitute the column purification from the original Abcam ChIP kit protocol with purification using AMPure XP beads (**see procedure step 35**). Generally, we enriched 10 % of input DNA by histone H3 and only 1 % by modified histone H3 antibody using 5- or 10-grams buds to performed ChIP experiment for 3 antibodies (**Table 2**). The amount of enriched-DNA from the modified histone H3 was considered too low for protocol validation using quantitative polymerase chain reaction (ChIP-qPCR) or conventional library construction for several reasons. First, our qPCR titration experiment showed that the lowest DNA concentration that can be detected by the qPCR machine should be no less than 0.1 ng/ µL (**Supplementary Information Table S1**). Second, there was no available positive control DNA target region for native- or modified-histone H3 in grapevine that could be used for ChIP protocol validation by qPCR. Lastly, library construction results were highly variable when DNA template was less than five ng. Based on these results, we suggest that 10 grams of buds (± 400 buds) may sufficient for one ChIP experiment only, i.e. immunoprecipitation of one protein of interest (e.g. modified histone H3) and one control (e.g. histone H3 or IgG).

**Table 1.** The yield of DNA using three different purification method.

| Purification method | DNA conc.*<br>(ng/μL) | DNA yield<br>(μg/grm) |
| --- | --- | --- |
| Abcam kit column purification | 0.71 | 0.14 |
| Phenol/Chloroform/Isoamyl alcohol | 6.56 | 1.31 |
| AMPure XP beads | 31.53 | 6.31 |

Note: DNA concentration was measured using Qubit fluorometer

**Table 2.** The average yield of input and ChIP-enriched DNA resulted from ChIP experiment using 5 and 10 grams of bud tissue for chromatin extraction (n = 3)

| Sample name | 5 grams | 10 grams |
| --- | --- | --- |
|  | Yield (ng) | Yield (ng) |
| MH_input | 274.8 | 398.7 |
| MH_histone H3 | 32.0 | 29.9 |
| MH_H3K4me3 | 1.1 | 3.2 |
| MH_H3K27me3 | 6.2 | 3.2 |
| MW_input | 305.6 | 412.2 |
| MW_histone H3 | 28.2 | 36.8 |
| MW_H3K4me3 | 1.3 | 2.8 |
| MW_H3K27me3 | 1.9 | 3.5 |
| May_input | 244.7 | 305.3 |
| May_histone H3 | 19.6 | 24.5 |
| May_H3K4me3 | 1.1 | 2.4 |
| May_H3K27me3 | 2.8 | 2.9 |
| Aug_input | 264.4 | 285.7 |
| Aug_histone H3 | 16.7 | 13.4 |
| Aug_H3K4me3 | 0.9 | 2.5 |
| Aug_H3K27me3 | 1.4 | 2.3 |

Abbreviations: MH, March H<sub>2</sub>CN<sub>2</sub> treated buds, MW, March water treated buds.
Note: On May and August buds were only treated with water.

### Antibody validation

Antibody recognition in grapevine buds was confirmed by Western blot analysis of grapevine buds nuclear extract recognising a ∼ 17 kDa band corresponding to predicted molecular weight of histone H3 and H3K4me3. The ImageJ software was used to estimate the signal intensity produced by each antibody (data not shown).

Immunoblot against anti-histone H3 showed detection limit of the antibody is around 40 ng and 200 µg nuclear extract containing a little less than 320 ng histone H3 protein (**Figure 8, panel 1)**. Anti-H3K4me3 passed the test showing absence of signal against 40 ng recombinant histone H3 protein (unmodified), and nuclear signal was about the half of nuclear signal produced against histone H3 antibody (**Figure 8, panel 2**). A false-positive signal observed against 320 ng recombinant histone H3 protein was observed; however, the intensity of the signal is no more than one-tenth the nuclear signal. No signal was observed in the nuclear extract tested against the anti-H3K27me3. We recognise that the lack of signal did not definitively indicate failure of the antibody, as this may result from low abundance of the modified histone in the tissue used for this test (**Figure 8, panel 3**).

**Figure 8.**
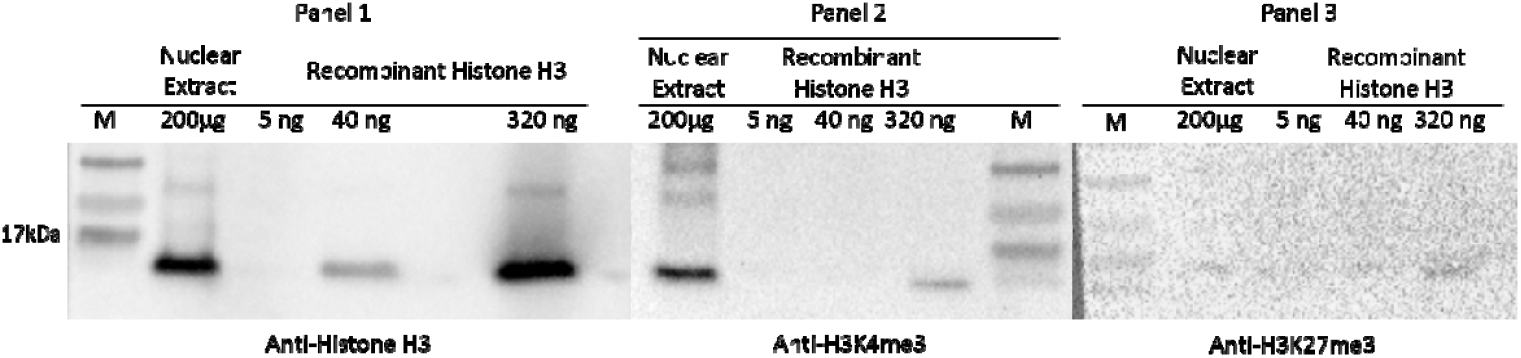
Representative western blotting assay for ChIP-antibody validation. Three antibodies used in ChIP assay were used for immunoblotting against nuclear extract prepared from grapevine buds and recombinant histone H3 at the concentration indicated in the image above. All antibodies were considered to pass validation test with detection of histone H3 protein and negative signal in H3K4me3 and H3K27me3 protein at 40 ng.

### Histone H3 occupancy

We generated an average 40 million 150 bp paired end reads from one replicate each of the histone H3-enriched and input DNA libraries of water-treated March (3W), May (5W), August (8W), and H_2_CN_2_-treated March buds (3H) buds. Although statistical comparisons cannot be made, it is worthwhile describing the trends. About 90 % of reads remained following trimming and were mapped uniquely to grapevine reference genome (**Supplementary Information Table S2**). Here, we showed a peak binding distribution of histone H3 at regions 4000 bp up- and down-stream of TSS in each condition. The highest occupancy was observed at the genic (exon, intron, or intergenic) region (**Figure 9**). ChIP peak calling analysis identified a different number of peaks at each condition, with the highest found in the May and H_2_CN_2_-treated March conditions, and the lowest in the water-treated March and August conditions (**Figure 9**).

**Figure 9.**
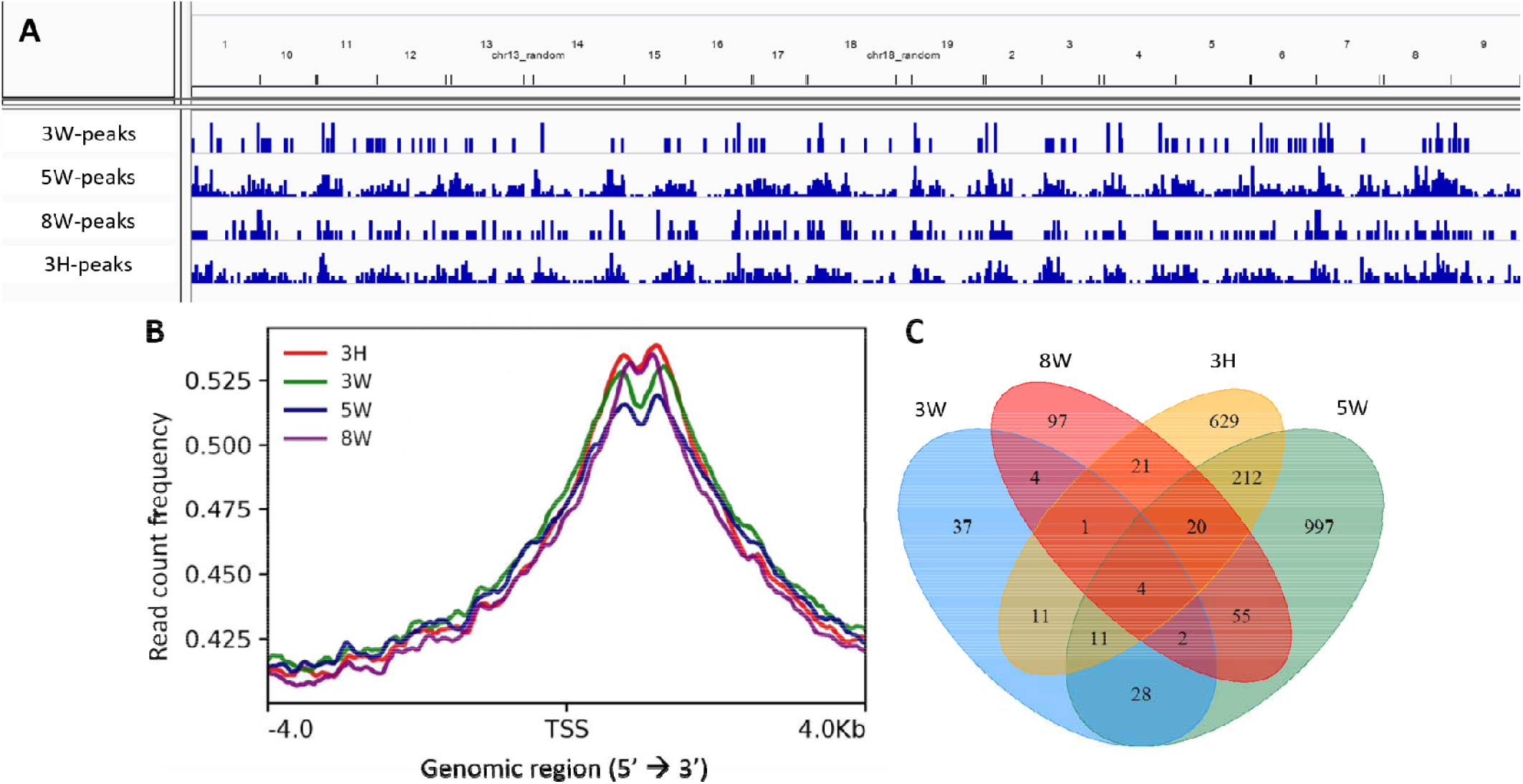
Chromatin immunoprecipitated-DNA peak analysis. (**A**) Distribution of histone H3 peaks along *Vitis vinifera* genome at each condition. (**B**) The average profile of ChIP peak binding at the transcription start site (TSS) region showing read count frequency range from −4000 to 4000 bp. (**C**)The Venn diagram of genes identified downstream TSS from buds collected in March, May, August treated with water and March buds treated with H_2_CN_2_.

A comparison between nucleosome occupancy and gene expression in *Arabidopsis* showed that genes with higher transcript abundance tend to be relatively unoccupied by nucleosomes at the promoter area, but relatively enriched in the genic region immediately downstream of the TSS (Valouev et al., 2011; Li et al., 2014b). We then restricted the Venn analysis and gene ontology (GO) enrichment to gene identifiers that were only enriched at the genic region (not the promoter region). The Venn diagram analysis shows that only few genes were commonly identified across samples, except for the May condition (5W) and March H_2_CN_2_ treatment (3H), with 247 common genes (**Figure 9**).The GO enrichment for gene identifiers at each condition is summarised using Treemap generated by REVIGO (**Supplementary Information Figure S1**). Relatively few biological processes were enriched in water-treated March and August condition buds by comparison with the May condition and buds treated with H_2_CN_2_. Categories related with meristem developmental state were enriched in water-treated March and May condition represented by embryonic morphogenesis (GO:0048598) in March and post-embryonic development (GO:0009791) in May. Meanwhile, the response to cold (GO:0009409) category was enriched coincident with prolonged exposure to cold in the August condition. Enrichment of categories related with cell growth (GO:0016049) and cell differentiation (GO:0030154) was seen in H_2_CN_2_-treated buds (**Supplementary Information Table S4**), suggesting regulation of growth at multiple levels. Further, we performed GO enrichment for the common gene identifiers found in May and H_2_CN_2_-treated buds (**Figure 10, Supplementary Information Table S5**). The results showed enrichment of categories related with response to starvation (GO:0042594), post-embryonic development (GO:0009791), and the regulation of phase transitions from vegetative to reproductive (GO:0048510) in both conditions. The genes associated with the enriched category were found to be involved in autophagy, flowering time, reactive oxygen species detoxification, sugar signalling, ABA-mediated signalling, and pleiotropic responses (**Table 3**).

**Table 3.**
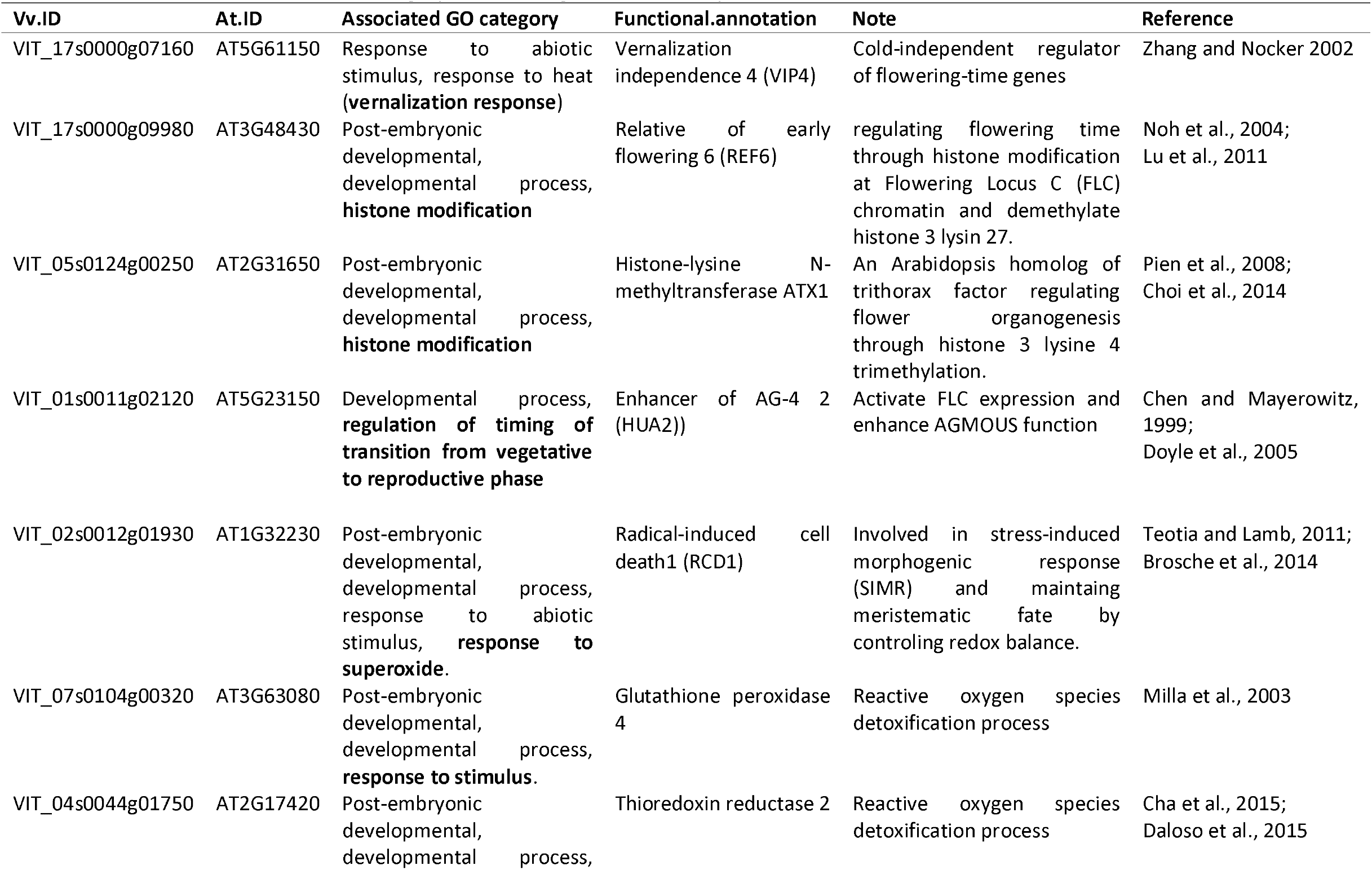
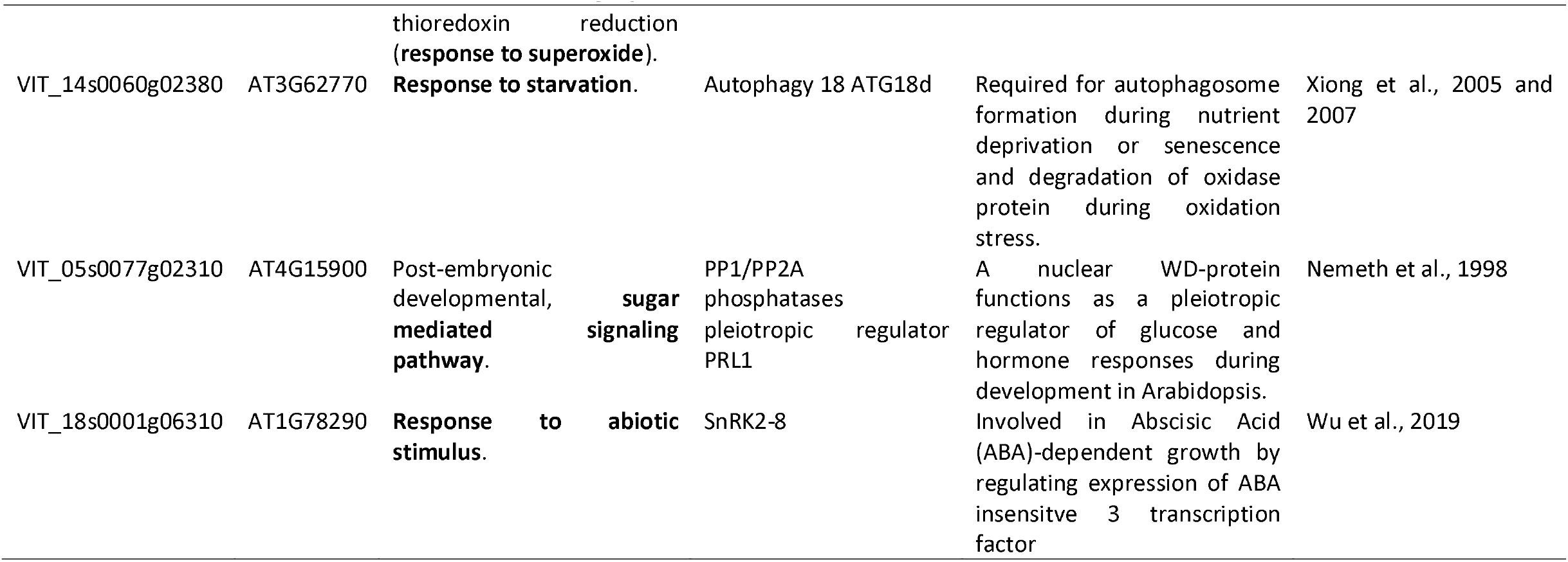
Gene associated with enriched category of common gene found in May and H_2_CN_2_-treated buds.

**Figure 10.**
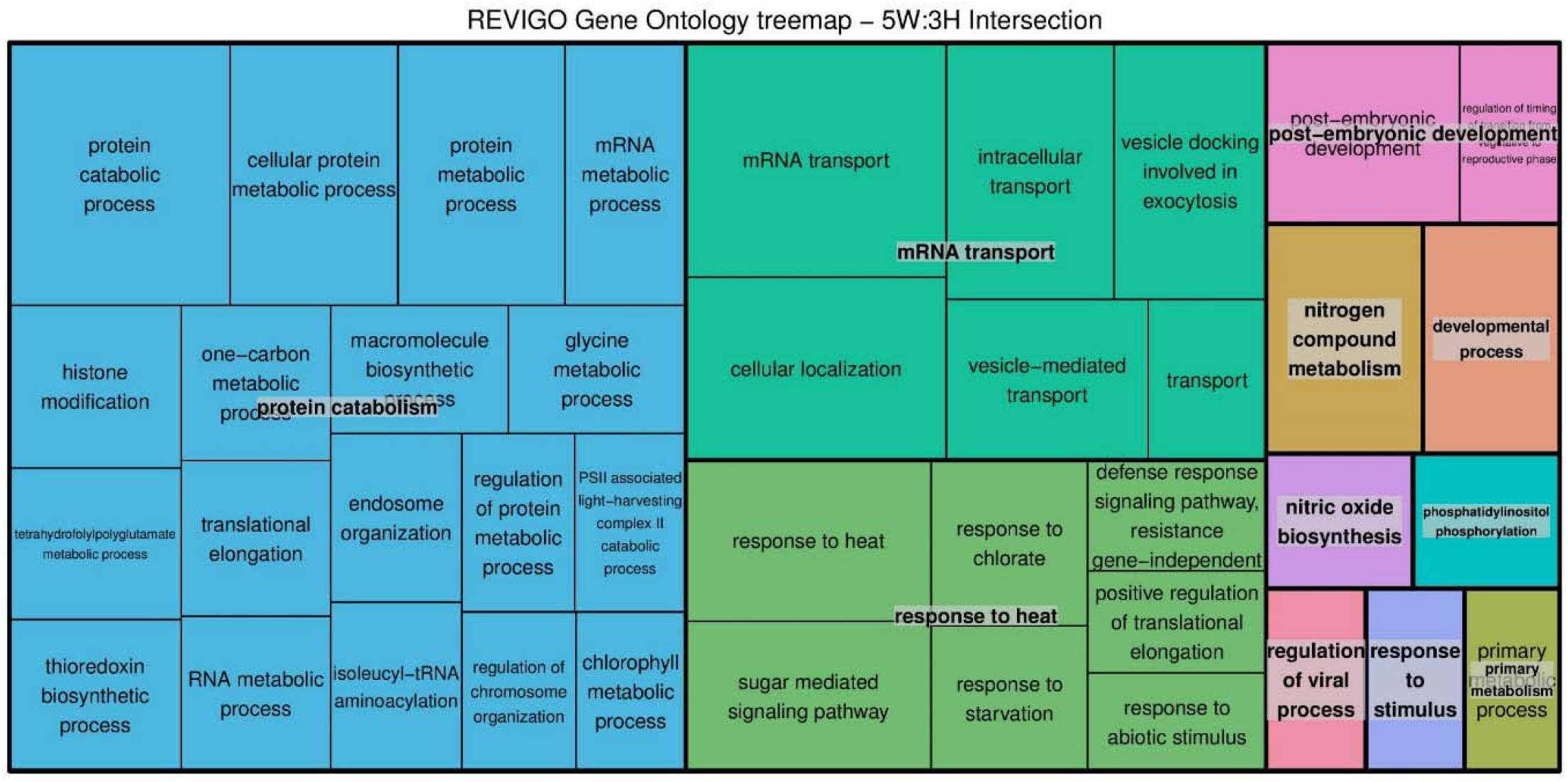
Functional category enrichment of genes associated with histone H3 commonly found in May, water-treated buds and March, H_2_CN_2_-treated buds. The highly redundant list of gene ontology (GO) terms is summarised and visualised using the TreeMap of REVIGO. The TreeMap view show two hierarchical level of GO terms. First, the semantically similar terms are grouped it to a representative subset (a non-redundant terms) visualised in a single rectangular. Second, the representative subsets are then clustered into a more general terms (printed over the box graphic) visualised by colours. Box size reflect the *p*-value of each non-redundant term.

## Discussion

### Optimisation conditions

#### Plant Material

The amount of tissue used in ChIP experiment with plant tissue varies depending on tissue type. Several early studies used 100 grams tissue per ChIP experiment (Ascenzi and Gantt, 1999; Chua et al., 2001) but recent improvements have enabled efficient ChIP with 1-5 grams, or 1×10^5^ purified nuclei (Gendrel et al., 2005; Deal and Henikoff, 2011). The axillary buds of grapevine are heterogeneous organs consisting of multiple vegetative and reproductive meristems and leaves, covered in trichome hairs (**Figure 11**). Considering that the buds consist of very little green tissue, we expected that nuclear density may be low. Our experiment demonstrated that 400 buds (± 10 grams) was only enough for ChIP experiment using one protein of interest (e.g. modified histone H3) and one control (e.g. histone H3 or IgG).

**Figure 11.**
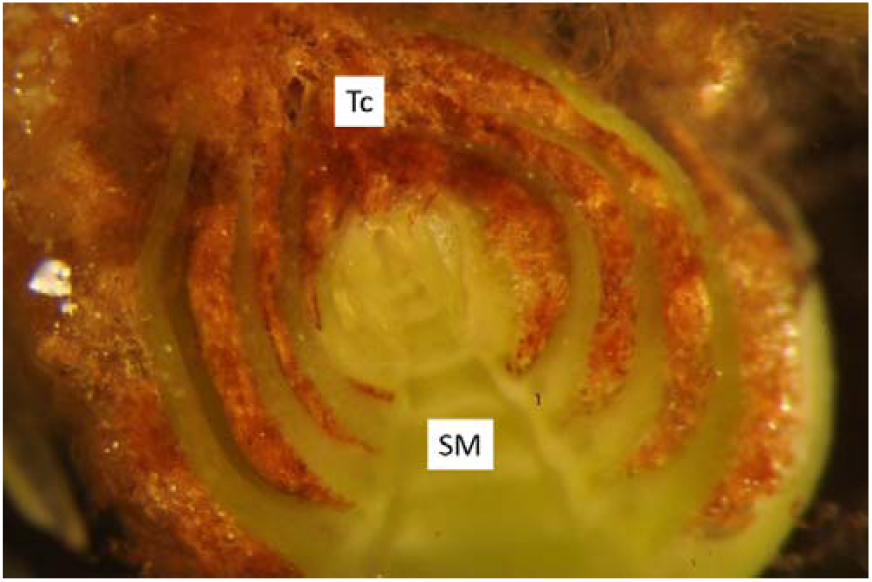
Anatomy of primary meristem the grapevine axillary bud. Trichome (**Tc**) hairs are shown as the brown-colour structures which surrounds the green tissue (**SM**, shoot meristem) of the axillary bud.

#### Crosslinking

Optimising the incubation conditions for crosslinking is crucial for successful and efficient crosslinking (Orlando, 2000). A short incubation duration for crosslinking is preferred in a ChIP experiment. Established protocol with yeast (Shivaswamy and Iyer, 2007), alga (Strenkert et al., 2011), animal (Browne et al., 2014) or plant (Li et al., 2014a) cells usually apply 10-30 minutes incubation for crosslinking procedure. However, the hair-like structures inside buds create air spaces which could impede penetration of the crosslinking solution. The application of a vacuum cycle procedure, was used here to change the pressure around the buds and remove entrapped air, thus allowing more efficient infiltration (Li et al., 2014a; Clode, 2015). To test the efficiency of our vacuum infiltration technique, we performed de-crosslinking followed by DNA extraction using the phenol:chloroform: isoamyl-alcohol (PCI) method. An optimal crosslinking must allow reversal of the process by heating (Das et al., 2004) and should result a maximum recovery of DNA by the PCI extraction (Haring et al., 2007; Ricardi et al., 2010). We conclude that the crosslinking duration should be limited to a maximum of 30 minutes and suggest performing crosslinking in batches, i.e. 15 minutes for excising buds from the canes followed by 15 minutes crosslinking.

#### Chromatin extraction

In lignified tissues, the presence and composition of secondary metabolites creates a requirement to optimise extraction conditions, particularly the composition of the homogenisation buffer and presence and concentration of detergent used for cell lysis (Li et al., 2014a). A powerful homogeniser such as the ULTRA-TURRAX (IKA, Germany) is also strongly recommended to improve tissue homogenisation. Moreover, polyvinylpyrrolidone (PVP) has been used routinely in nuclei acid extraction from tissue with high polyphenol content (Lodhi et al., 1994; Porebski et al., 1997). Secondary metabolites, such as polyphenols and tannins, can bind to DNA upon cell lysis and contaminated DNA may present problem for downstream analysis, such as DNA library construction for sequencing. The PVP binds polyphenols through hydrogen bonding and can then be removed from tissue homogenate by discarding the supernatant containing PVP-polyphenols after centrifugation step (John, 1992). There are also several considerations in the choice and amount of detergent. Typically, an anionic detergent such as sodium docecyl sulfate (SDS) is used, however while concentrations > 0.1 % SDS (w/v) will improve nuclear isolation, this may disrupt the antibody-antigen interaction due to protein denaturation (Privé, 2007). Moreover, high concentrations of ionic detergent tend to result in formation of precipitates at low temperature, risking inefficient cell lysis and co-precipitation with the DNA (Linke, 2009). Two concentration of SDS commonly used in ChIP assays were tested here, i.e. 0.1 % and 1 %, to determine the optimum condition resulting in the highest yield of DNA for immunoprecipitation. Also, we tested 0.1 % sarkosyl, a milder anionic detergent which is structurally similar to SDS but remains soluble under low temperature, as a comparison to the widely use SDS (Linke, 2009). Our result show that lower detergent concentration, both ionic and anionic, resulting a low DNA yield (**Figure 5, lane 1, 3, and 5**). However, the result was improved after sonication was applied for several minutes.

#### DNA fragmentation

The most common procedures to shear DNA for ChIP assay is by sonication (Orlando, 1997 and 2000) or micrococcal nuclease treatment (O’Neil et al., 2003); the former method is mainly used for crosslinked ChIP experiment. Ideally, DNA is sheared into small fragment range from 200 to 600 bp (Park, 2009). Sonication is highly variable and difficult to optimise. A titration approach is commonly required to find the best sonication duration and settings. By considering this, we then performed a test to determine the sonication duration that will produce the desired fragment size. Here, we use S220 Focused-Ultrasonicator (Covaris, USA) and followed manufacture recommendation to generate homogenously distributed ∼300 bp DNA fragment, i.e. 5 % Duty Cycle, 4 intensity, 140 W peak incident power, 200 cycles per burst. We then tested three sonication duration, i.e. 6, 8 and 10 minutes. Fragmented DNA was then analysed using TapeStation^®^ (Agilent, Australia) and quantified using Qubit (Thermo Fischer Scientific, Australia) as both methods provide a more sensitive and accurate measurement comparatively to measurement using agarose gel or nanodrop respectively (Simbolo et al., 2013). The sonication step served two purposes in our protocol, i.e. improve cell lysis and DNA fragmentation. Aggregated nuclei are a common problem when isolating nuclei from tissue with high tannic acid content (Loureiro et al., 2006) and clumping nuclei will also reduce efficiency of DNA fragmentation (Arrigoni et al., 2015). Development of a standard ChIP protocol using animal cells also demonstrates that mild sonication can help to separate clumping cells which then improve cell lysis process and increased DNA yield (Arrigoni et al., 2015). In agreement with this report, our result showed that the use of high detergent concentration for cell lysis could be avoided using our sonication settings. In addition to improve cell lysis, our sonication setting was found to be affected long DNA more than short DNA. Library construction may increase bias toward short DNA fragments due to size selection during library construction. Although 10 minutes sonication was sufficient to shear grapevine chromatin into a suitable size for sequencing (usually within 150-300 bp range), we suggest to apply 12 minutes sonication in order to obtained a higher amount of DNA fragment within the 150-300 bp range.

#### Antibody validation

A specific antibody with high affinity to the protein of interest is a prerequisite for a successful ChIP experiment (Kungulovski et al., 2015). Antibodies are common tools to study many biological processes; however, they may also cause problems (Saper and Sawchenko, 2003; Baker, 2015a). Common problems are (1) recognition of non-target protein due to antibody cross-reactivity, (2) non-reproducible results due batch-to-batch variation of antibody, and (3) unsuitable application, for example antibodies that work for western blotting may not suitable for immunoprecipitation (Baker, 2015a). It is imperative to characterise and validate the antibody of choice before commencing an experiment (Schumacher and Seitz, 2016; Gautron, 2019). Egelhoffer et al. (2011) tested 246 ChIP-grade antibodies and found many of these antibodies were either non-specific or unsuitable for ChIP. In order to address this issue, we performed antibody assessment to validate the ChIP antibody that was used in our experiment. We chose antibodies for histone H3, H3K4me3, and H3K27me3 on the basis of existing public data on the specificity, in order to meet at least one of the selection criteria. The antibodies chosen had been shown to specifically recognise the antigen in HeLa cells by the manufacture, in various human or mouse tissue by the ENCODE project and used in ChIP analysis in barley (Baker et al., 2015b). Recombinant histone H3 and nuclear extract of grapevine buds were tested against anti-histone H3, anti-H3K4me3, and anti-H3K27me3. Criteria for an antibody to “pass” specificity by western blotting was adopted from Egelhoffer et al. (2011), i.e. the tested antibody should produce at least 50 % signal compare to the total nuclear signal and ten-times higher than any unspecific signal.

### ChIP-sequencing and Histone H3 occupancy

The outcome from the ChIP experiment is fragments of DNA that specifically interact with the protein of interest. Identification of the DNA sequence following the immunoprecipitation can be done by polymerase chain reaction (ChIP-PCR) or quantitative PCR (ChIP-qPCR), microarray (ChIP-chip), and high-throughput sequencing (ChIP-seq). The endpoint PCR or qPCR is the most widely and routine identification technique use in ChIP. The pitfall of this technique is that it requires prior knowledge of regions associated with the protein tested. Rapid improvement of genome-wide assays using microarray or high-throughput sequencing, provide an alternative DNA assay for species such as grapevine; in which knowledge about the region occupied by histone H3 or modified histone H3 is not available. Several reviews outline the superiority of sequencing over microarray for several reasons, such as higher genome coverage including the repeated sequence and low noise to signal ratio which commonly found in microarray analysis (Schones and Zhao, 2008; Park, 2009; Furey, 2012). In this study, we performed a ChIP-seq analysis of the histone H3 to evaluate our ChIP protocol. We also compare and explore the histone H3 occupancy along grapevine bud chromatin using dormant buds harvested at three different time point. Nevertheless, differential regulation of histone H3 is beyond the scope of this protocol.

Nucleosome (histone octamer) occupancy and positioning have been suggested to play important roles in regulating gene expression and many additional DNA-related process (Struhl and Segal, 2013). Studies of nucleosome occupancy and positioning in animals, yeast, and plant cells have demonstrated a bias in nucleosome occupancy positioning towards regions proximal to the transcription start site (TSS) (Mavrich et al., 2008; Schones et al., 2008; Lee et al., 2017; Zhang et al., 2019). Furthermore, genome-wide nucleosome occupancy studies in yeast, mammalian and plant systems showing that the genomic sequence of nucleosome is mostly depleted in the promoter or transcription termination sites (Field et al., 2018; Fenouil et al., 2012; Liu et al., 2015). In yeast, nucleosome depletion was found in the homopolymers of deoxyadenosine nucleotides (poly (dA:dT) tracts) regions, suggesting that the structure of poly (dA:dT) tracts may be resistant to the bending and twisting deformation required to wrap DNA around nucleosomes (Field et al., 2008; Segal and Widom, 2009 and the reference therein). On the contrary, in mammalian and plant tissues, promoter regions are mostly GC-rich, hence the nucleosome depletion is tightly associated with CpG islands (Fenouil et al., 2012; Liu et al., 2015). Our result showing a similar pattern of histone H3 occupancy with those previously reported in study with the histone octamer, i.e. higher preference occupation at down-stream TSS region. Functional category analysis of gene identifier at the genic region showed enrichment of process related with meristem development and response to environment condition at the time of sampling, e.g cell cycle activities. Differential expression and abundance of histone H3 was reported to correlate well with DNA synthesis and cell cycle activities, showing highest abundance during early embryogenesis in *Drosophilla* (Shindo and Amodeo, 2019), or in cycling cells of plant meristems (Kaparos et al., 1992; Terada et al., 1993; Sano and Tanaka, 2005) and at low abundance in quiescent apical buds (Singh et al., 2009). Annotation of the DNA associated with the histone H3 peaks identified 129, 1691, 291, and 1207 genes for the 3W, 5W, 8W and 3H conditions (**Supplementary Information Table S3**).

## Conclusion

We describe the systematic optimisation of detail chromatin immunoprecipitation protocol for grapevine bud samples. The protocol was developed from ChIP protocol for woody tissue published by Li et al. (2014a) and then modified according to optimisation results that we performed at each step of the ChIP protocol; this included the amount of starting material, crosslinking method, chromatin extraction condition, chromatin shearing duration, validation of antibody, and DNA purification method. Identification of histone H3 enriched DNA by sequencing, provided an example for the potential use of this protocol to study the post-translational modification of histone H3 in the buds of grapevine. Comparing the results from nucleosome occupancy in yeast, human, and Arabidopsis we validated our ChIP experimental data.

## Data availability statement

All datasets generated for this study are included in the article/ supplementary material.

## Author contribution

M.J.C. conceived and supervised the project. D.H. is responsible for data curation, analysis, and investigation. Optimisation of the ChIP procedure was performed by D.H. in collaboration with J.C. and R.L. T.C.* performed the sequencing and data processing. D.H. wrote the manuscript with constructive comment from M.C. and T.C.* All authors contributed to the article and approved the submitted version. *T.C. deceased prior to submission but after approving the submitted manuscript.

## Acknowledgements

We express deep sadness at the passing of Tinashe Chabikwa and sincere condolences to his family. We are thankful to Keith Mugford of Moss Wood Wines for enduring support and access to plant material at often inconvenient times. We are also very grateful to the team of the Centre for Microscopy, Characterisation and Analysis of the University of Western Australia for technical guidance.

